# Birth of new protein folds and functions in the virome

**DOI:** 10.1101/2024.01.22.576744

**Authors:** Jason Nomburg, Nathan Price, Jennifer A. Doudna

## Abstract

Rapid virus evolution generates proteins essential to infectivity and replication but with unknown function due to extreme sequence divergence^1^. Using a database of 67,715 newly predicted protein structures from 4,463 eukaryotic viral species, we found that 62% of viral proteins are evolutionarily young and lack homologs in the Alphafold database^2,3^. Among the 38% of more ancient viral proteins, many have non-viral structural homologs that revealed surprising similarities between human pathogens and their eukaryotic hosts. Structural comparisons suggested putative functions for >25% of unannotated viral proteins, including those with roles in the evasion of innate immunity. In particular, RNA ligase T- (ligT) like phosphodiesterases were found to resemble phage-encoded proteins that hydrolyze the host immune-activating cyclic dinucleotides 3’3’ and 2’3’ cyclic G-A monophosphate (cGAMP). Experimental analysis showed that ligT homologs encoded by avian poxviruses likewise hydrolyze 2’3’ cGAMP, showing that ligT-mediated targeting of cGAMP is an evolutionarily conserved mechanism of immune evasion present in both bacteriophage and eukaryotic viruses. Together, the viral protein structural database and analytics presented here afford new opportunities to identify mechanisms of virus-host interactions that are common across the virome.

## Introduction

Viral proteins carry out functions critical for infection. Some proteins or their component domains are widely conserved within and across viral families, including between viruses of distinct Baltimore classifications^4^ and in viruses that infect different kingdoms of life^5–8^. These include “viral hallmark genes”, such as the jelly-roll folds of viral capsid proteins and folds related to RNA- and DNA-directed RNA polymerases ^4,9^. However, a major challenge to understanding viral infection mechanisms and evolution is the large percentage of viral proteins with unknown function. Sequence homology between viral proteins and other viral or non-viral proteins can sometimes suggest protein function, but the rapid pace of viral evolution and *de novo* emergence of genes generate many proteins without annotated sequence homologs. This creates a pressing need for alternative approaches to identify protein homologs.

Viral proteins are highly divergent even within the same family, limiting the utility of sequence-based homology searches when amino acid identity falls below ∼30%^10–13^. By contrast, horizontal gene transfer among viruses and between viruses and cells creates structural relationships that can inform about protein function if they can be detected^3,14–17^. However, viral proteins have limited representation among experimentally determined structures in the Protein Databank (PDB) and they are absent from the predicted protein structures in the AlphaFold Database^2,18,19^.

To address this gap and develop a means to systematically predict viral protein functions, we generated a database of predicted structures from 67,715 proteins encoded by 4,463 species of eukaryotic viruses. We clustered these proteins by both sequence and structure, generating 5,770 multi-member and 12,422 singleton clusters. Structural homology searches greatly expanded the taxonomic diversity of protein clusters, revealing putative protein function by connecting unannotated viral proteins with annotated homologs. Structural comparisons between viral and non-viral proteins identified potential functions encoded by human pathogens, including previously undetected synergies between anti-immunity activities. In particular, ligT-like phosphodiesterases (PDEs) emerged from this analysis as a widespread class of enzymes conserved across the bacterial and eukaryotic virome. Conservation evident within our viral protein structure database, together with enzymatic activities validated in cell-based experiments, reveal an ancient and fundamental role of these proteins in viral anti-immunity pathways.

### The structural proteome of eukaryotic viruses

To analyze the diversity of protein structures present in eukaryotic viruses, we used Colabfold^20^ to predict the structures of 67,715 proteins from eukaryotic viruses included in RefSeq. We then implemented a two-step approach to cluster them, using both sequence-based and structure-based clustering (Fig. 1A)^3^. We used Mmseqs2^21^ to cluster protein sequences to 70% coverage and 20% identity, resulting in 21,913 sequence clusters. Next, we leveraged the alignment speed of Foldseek^22^ to conduct structural alignments between a single representative of each sequence cluster and filtered alignments to keep those with at least 70% alignment coverage, a TMscore of at least 0.4, and an *E*-value lower than 0.001. The resultant structural alignments had a median TMscore of 0.52 (Supplementary Fig. 1A), reflecting robust structural similarity^23^. The 70% alignment coverage threshold enriches clusters for members that are similar across the majority of their protein sequence. Cumulatively, this resulted in 18,192 protein clusters, of which 12,422 have a single member (Supplementary Fig. 1B). This dataset includes a large diversity of viruses, including 4,463 species from 132 different viral families (Fig. 1B, C). Clusters are structurally consistent, as implementing DALI^24^ to align cluster representatives to each member for clusters with at least 100 members yields a median cluster-average DALI Z score of 13.1 (Supplementary Fig. 1C). DALI Z-scores above 8 indicate two proteins are likely homologous^25^.

**Fig. 1:**
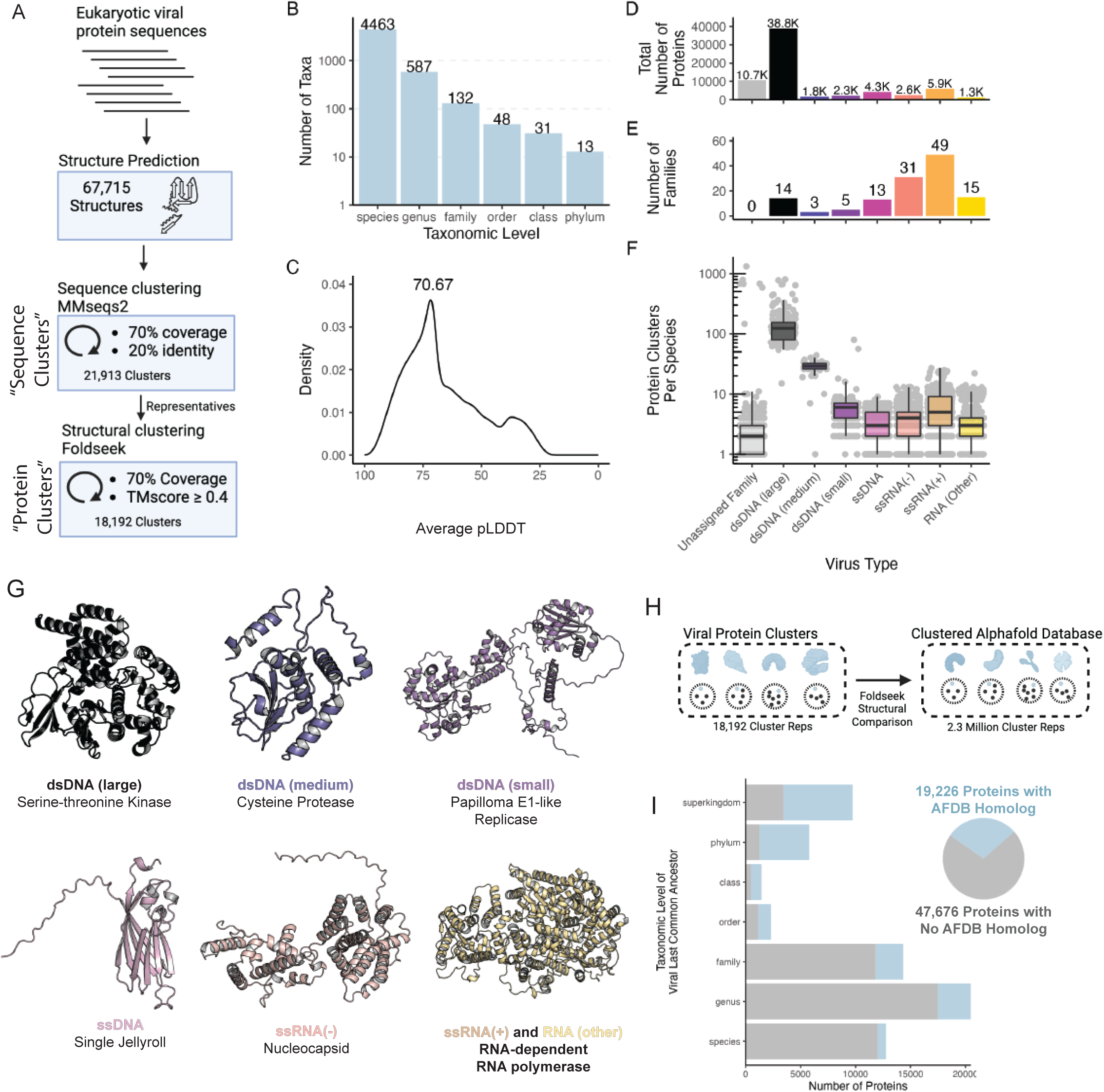
The structural proteome of eukaryotic viruses. (**A**) Pipeline for protein clustering. First, proteins from viruses in RefSeq with an annotated eukaryotic host were folded using Colabfold, leading to generation of 67,715 predicted structures. Protein sequences were clustered to 70% coverage and 20% identity, resulting in 21,913 sequence clusters. The predicted structures of the representatives of each cluster were then aligned and clustered together with a requirement of 70% coverage across the structural alignment and a TMscore >= 0.4. This resulted in a final set of 18,192 clusters. (**B**) Taxonomic distribution of the dataset. Each column indicates the number of taxa present in the dataset. (**C**) Distribution of the average pLDDT of all structures in the dataset. (**D-F**) Viral families were classified by genome type into one of eight categories. For each category, the total number of proteins (D), number of viral families (E), and number of protein clusters per species (F) are indicated. (**G**) Protein structures represent the protein cluster that is encoded by the highest number of viral families of each genome type. (**H**) Foldseek was used to align a single representative protein from each viral protein cluster against a cluster representative from 2.3 million clusters generated from the Alphafold database. (**I**) (Left) The taxonomic level of the last common ancestor of each viral protein cluster was determined. For example, if a protein cluster is encoded by viruses from three different families from two different orders but the same class, they are placed in the class row. Proteins were split by this taxonomic level on the Y axis, with protein counts on the X axis. Blue color indicates the proteins belong to a cluster with a homolog in the AFDB, while gray indicates the proteins belong to a cluster without a homolog in the AFDB. (Right) A pie chart indicating the total number of proteins that belong to clusters whose representatives aligned to the AFDB (blue) or did not align (grey).

We investigated how well this database represents viral diversity, and if it reconstitutes core viral hallmark genes. We grouped viral families into viral genome types based on their Baltimore classes^26^ with slight modifications – DNA viruses were split into large, medium, and small groupings based on their average genome length, while RNA viruses without single-stranded positive- or negative- sense genomes were grouped into the RNA (Other) category. Large double-stranded DNA (dsDNA) viruses have the most protein clusters per species and, despite constituting only 14 of the 132 viral families in the dataset, account for the majority of viral proteins (Fig. 1D, F). As expected, protein cluster count correlates strongly with genome size (Supplementary Fig. 1D). With their larger genomes, dsDNA viruses have the capacity to encode more auxiliary genes without sacrificing genome stability. RNA viruses make up a large fraction of the families present in the dataset, but a smaller fraction of the total proteins (Fig. 1E, F). Structural homology between viral families with a similar genome type is common, with large dsDNA viruses sharing many protein folds (Supplementary Fig. 1E).

As expected, the predominant protein clusters in the dataset as a whole (Supplementary Fig. 1F) and within each genome type (Fig. 1G) are largely involved in fundamental aspects of the viral life cycle. These include the single jellyroll fold, which comprises viral capsids and is present in viruses of all genome types. The double jellyroll fold also comprises viral capsids, although it is restricted to dsDNA viruses^27^. RNA viral families often encode nucleocapsids, responsible for packaging of viral RNA, and RNA-dependent RNA polymerases (RdRPs) responsible for genome replication. While the RdRP is universally conserved in RNA viruses, it is split amongst multiple protein clusters due to variation in protein length. In contrast, small dsDNA viruses such as papillomaviruses and polyomaviruses encode a viral replicase with conserved origin-binding and helicase domains. Altogether, we find that our structural database successfully reconstitutes conserved viral proteins across diverse viral subtypes.

We next investigated the taxonomic distribution of viral protein clusters. We conducted structural alignments of viral protein cluster representatives against 2.3 million cluster representatives from the entire Alphafold Database (AFDB)^3^ (Fig. 1H). For each virus protein cluster, we determined the last common ancestor of viruses encoding a cluster member. We found that 29% of protein clusters are present in multiple viral families, the majority of which are present in the Alphafold Database, suggesting that they are evolutionarily ancient (Fig. 1I). In addition, we found that 62% of viral proteins (or 55% of proteins from non-singleton clusters) are restricted to a single viral family and lack homologs in the AFDB (Fig. 1I). This shows that viral evolution generates substantial numbers of novel proteins that are absent from current structure databases.

### Structural similarities between annotated and uncharacterized viral proteins

We investigated the ability of structural alignments to identify relationships not apparent from protein sequence alone. We found that many representatives of sequence clusters are structurally similar despite low sequence similarity (Fig. 2A). In fact, adding structural information to protein clustering efforts leads to more taxonomically diverse protein clusters, with significantly more viral families per cluster (Fig. 2B). This is especially important for finding homology between proteins from divergent viruses, resulting in a substantial increase in protein clusters encompassing proteins encoded by viruses of different genome types (Fig. 2C).

**Fig. 2:**
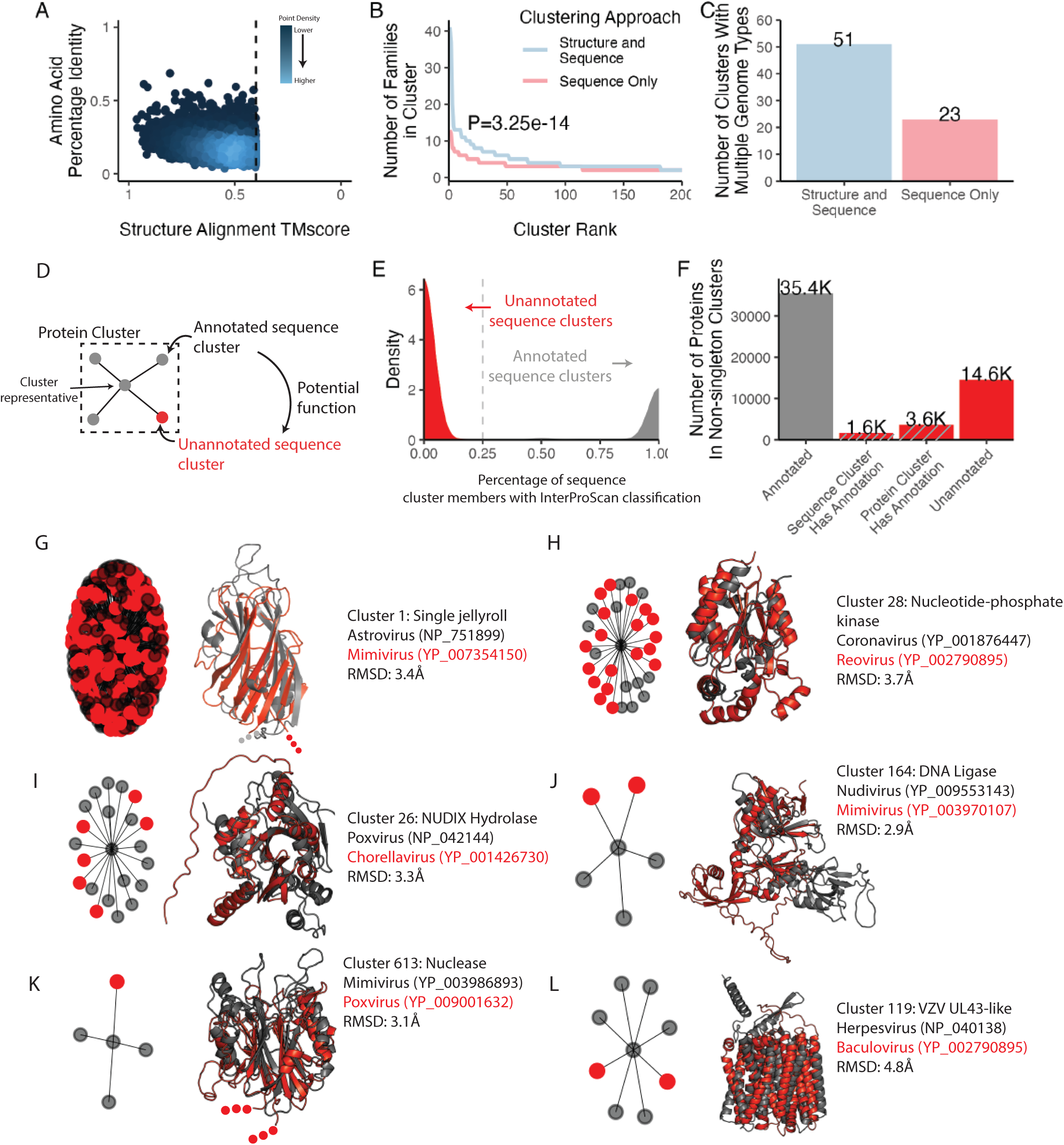
Structural alignments link annotated and unannotated sequence clusters. (**A**) Structural alignments identify homologs at low amino acid identity. Each dot indicates a single alignment between two sequence cluster representatives. The Y axis indicates the amino acid identity (from 1 to 0, with 1 indicating 100% of residues are identical). The X axis indicates the Foldseek TMscore, where a TMscore > 0.5 is highly confident and > 0.4 is somewhat confident. The color of each dot represents the density of points, with brighter colors indicating higher density. (**B**) Clusters generated with both structure and sequence information lead to significantly more viral families in each cluster when compared to clusters generated by sequence information alone (P<3.25e-14, wilcoxon rank sum test). Proteins were clustered by sequence and structure or by structure alone, and the top 200 clusters by number of members was plotted. The Y axis indicates the number of families in each cluster for each cluster on the X axis. (**C**) Incorporating structure information greatly increases detection of protein homologs present in different genome types. The bars indicate the number of clusters that contain proteins from viruses with different genome types when clusters are generated using structure and sequence or sequence only. (**D**) Protein clusters contain structurally-related sequence clusters. In some cases, sequence clusters that are poorly annotated (few members have InterProScan classifications) are in the same protein cluster as, and therefore structurally-similar to, sequence clusters that are well annotated. Thus, the annotations of annotated cluster members can provide information on the potential function of unannotated proteins that share structural similarity. (**E**) Protein clusters can be classified as unannotated or annotated based on the fraction of cluster members with an InterProScan classification. The percentage of sequence cluster members with an InterProScan classification was determined (X axis), and is plotted based on the density of sequence clusters (Y) with each percentage. Sequence clusters with < 25% of members that contain InterProScan classifications were considered unannotated sequence clusters. (**F**) Many unannotated proteins cluster via sequence or structure with proteins that are annotated. If a protein has an InterProScan classification, it is counted as Annotated. Proteins that are not classified but fall within either an annotated sequence cluster or a protein cluster that contains an annotated sequence cluster were also counted. (**G-L**) (Left) A network of sequence clusters that belong to each protein cluster, where nodes that are red are unannotated and those that are grey are annotated. The centroid is the protein cluster representative. (Right) Members of annotated and unannotated sequence clusters are highlighted, where the structure of an annotated protein (grey) is compared to the structure of an unannotated protein (red). The RMSD between the two structures is indicated.

We asked if structural alignments can link poorly-annotated sequence clusters with those that are more annotated (Fig. 2D). We used the sequence-based classifier InterProScan^28^ to assign all proteins Pfam^29^, CDD^30^, and TIGRFAM^31^ classifications. Sequence clusters contain almost entirely annotated or entirely unannotated members, resulting in a bimodal distribution of sequence clusters (Fig. 2E). Of the proteins in clusters with more than one member, over 25% of unannotated proteins are located in either an annotated sequence cluster or a protein cluster that contains an annotated sequence cluster (Fig. 2F).

Many protein clusters encompass a mixture of annotated and unannotated sequence clusters (Supplementary Fig. 2). We find that these connections between sequence clusters are useful to determine putative functions of poorly characterized proteins across the virome. For example, while the single jellyroll fold is the most abundant protein cluster, many members of this cluster are not correctly annotated (Fig. 2G). Many other protein clusters include both annotated and unannotated sequence clusters, including clusters encoding enzymes such as nucleotide-phosphate kinases (Fig. 2H), NUDIX Hydrolases (Fig. 2I), DNA ligases (Fig. 2J), and nucleases (Fig. 2K). One cluster of note includes members that resemble the UL43 family of late herpesvirus proteins (Fig. 2L), which will be discussed later. Together, these results demonstrate that large scale clustering based on sequence plus predicted structure enables functional inference of poorly characterized viral proteins.

### Structural alignments suggest functions of human pathogen proteins

Unlike nucleotide or protein sequence, structural features are often conserved over large evolutionary timescales. Thus, we investigated if alignment between predicted viral and non-viral protein structures can offer insight into the function of poorly-annotated proteins encoded by human pathogens. To do this, we used Foldseek to align our virus protein structure database with the initial release of the Alphafold Database, which contains over 300,000 proteins from 21 organisms across eukaryotes, bacteria, and archaea^2^ (Fig. 3A). This revealed pervasive structural homology between viral and non-viral proteins, with high structural similarity in the face of low amino acid identity (Fig. 3B).

**Fig. 3:**
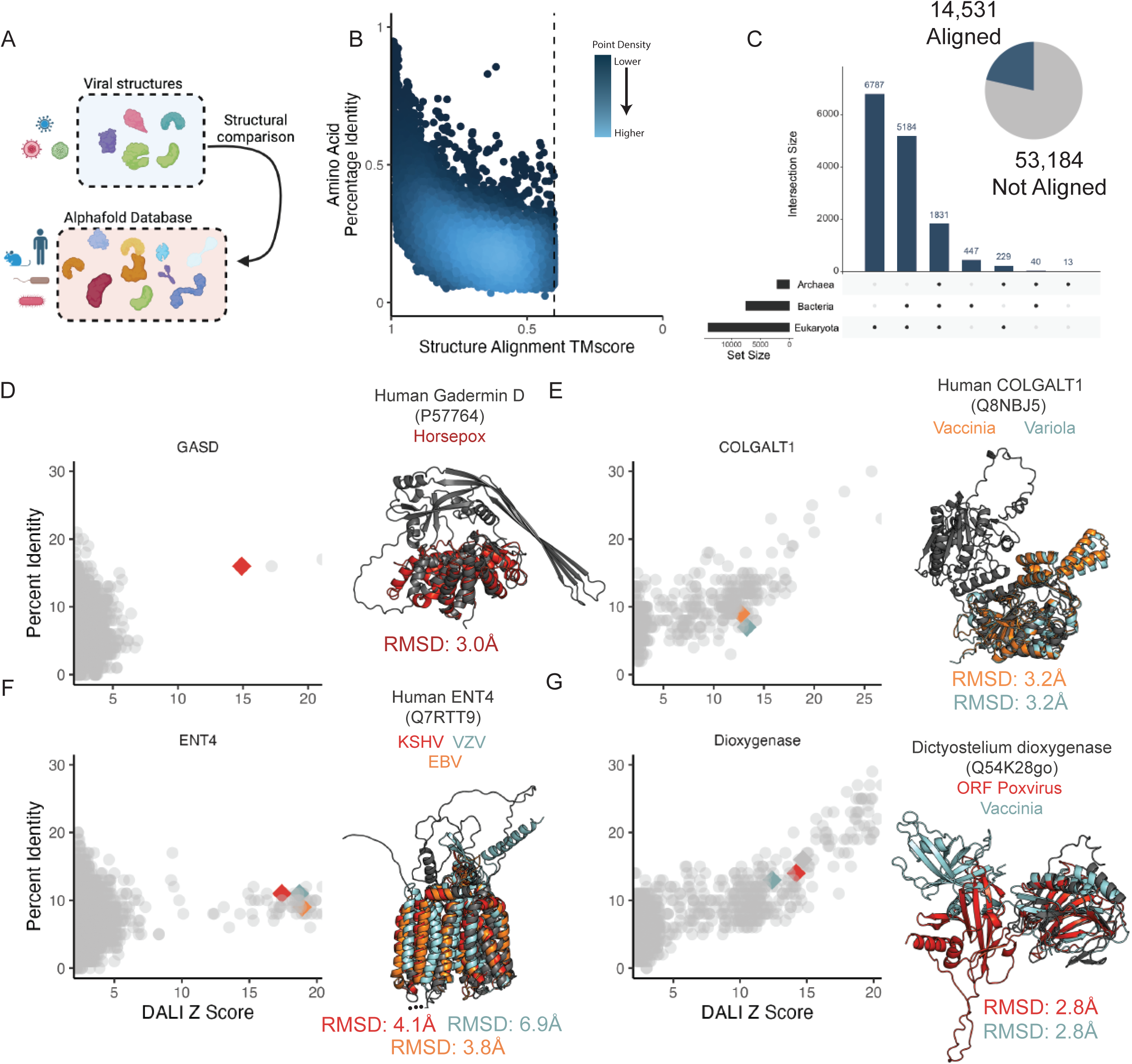
Structural homology across kingdoms of life reveals potential protein function. (**A**) Illustration of the approach. The database of viral protein predicted structures were aligned against the Alphafold database consisting of proteins from 48 organisms and including members of the bacterial, eukaryote, and archaeal superkingdoms. (**B**) A plot indicating the amino acid percentage identity (Y axis) and Foldeseek TMscore (X axis), where each point indicates a single alignment. For viral proteins with more than five alignments, the top five by TMscore are plotted. The color of each dot represents the density of points, with brighter colors indicating higher density. (**C**) (Right) A pie chart indicating the number of viral proteins that do or do not have an alignment against the Alphafold database. (Left) An UpSet plot indicating, for those viral proteins with alignments against the Alphafold database, the number that align against members of each superkingdom. (**D-G**). Specific non-viral hits from the Alphafold Foldseek search were aligned against the viral predicted structure database using DaliLite, and alignments against proteins from human pathogens were selected. (Left) The Y axis indicates the percentage amino acid identity, and the X axis indicates the Dali Z score. Each dot indicates a single alignment. Each point indicates an alignment, with the points corresponding to the proteins highlighted on the right as diamonds and colored consistently with their protein structures. (Right) The structure of the non-viral protein query is present in black. A superposition of selected protein clusters is shown, with the RMSD of each viral protein vs the non-viral protein indicated. Protein accessions are as follows: GASD: (Horsepox-ABH08278). COLGALT1: (Vaccinia-YP_232983; Variola-NP_042130). ENT4: (KSHV-YP_001129415; VZV-NP_040138; EBV-YP_401658). Dioxygenase: (ORF Poxvirus-NP_957891; Vaccinia-YP_232906)

Ultimately, 14,531 predicted viral proteins have an alignment to a member of the Alphafold database, with the majority of alignments being against proteins encoded by eukaryotes (Fig. 3C). These alignments include proteins that are unannotated but are encoded by human pathogens. To reduce rates of false negatives, we conducted a series of alignments using DALI^24^, which is slower than Foldseek but substantially more sensitive. First, we found that a set of proteins encoded by poxviruses are structurally similar to the auto-inhibitory domain of mammalian gasdermins (Fig. 3D)^32^. Similarly, several poxvirus proteins are structurally homologous to the human galactosyltransferase COLGALT1, thought to enable virus binding to surface glycosaminoglycans during viral entry (Fig. 3E)^33^. Next, we found that human herpesviruses proteins, including the protein BMRF2 from Epstein Barr herpesvirus (EBV) and Varicella zoster virus (VZV), share structural similarity with the human equilibrative nucleoside transporter ENT4 (Fig. 3F). EBV conducts substantial remodeling of host cell metabolism during viral infection^34^, and this finding suggests a potential metabolic role in addition to BMRF2 involvement in viral attachment^35^. In addition, transport of antiviral nucleoside analogues such as valacyclovir are mediated by nucleoside transporters^36,37^, raising questions about the interplay between this protein and valacyclovir during VZV infection. These proteins belong to a cluster of proteins similar to the UL43 family of late herpesvirus proteins, some of which are unannotated (Fig. 2J). In addition, we observed structural homology of Poxvirus C4-like proteins with eukaryotic dioxygenases (Fig. 3G). Vaccinia virus C4 is notable for antagonizing several innate immune pathways. C4 directly binds the pattern recognition receptor DNA-PK, blocking DNA binding and immune signaling through that pathway^38^. In addition, C4 inhibits NF-κB signaling downstream at or downstream of the IKK complex, but the mechanism of this inhibition is unknown^39^. Future studies are required to determine if its dioxygenase-like fold is involved in its innate immune antagonism. Altogether, these findings illustrate the ubiquity of structural homology between viral and non-viral proteins and show that this homology can be used to predict potential functions of poorly characterized viral proteins.

### Horizontal gene transfer creates taxonomically-diverse protein clusters

While we found that some protein clusters contain members encoded by viruses of different genome types, the evolutionary origin of such conservation is unclear. Many of these protein clusters are predominantly encoded by viruses of a single genome type but expressed in a small minority of viruses of a different genome type (Supplementary Fig. 3A). This observation is consistent with virus-virus or host-virus horizontal gene transfer.

To explore this possibility, we conducted Blast^40^ searches of sequence cluster representatives against viral- and non-viral protein databases and constructed phylogenetic trees of the top hits. We found that nucleoside-phosphate kinases in cluster 28 show a polyphyletic distribution with homologs in different viruses showing amino acid similarity to distinct sets of non-viral proteins (Supplementary Fig. 3B). There is a similar pattern with HrpA/B-like helicases in Cluster 55, with helicases in different viral families showing amino acid similarity to distinct sets of non-viral organisms (Supplementary Fig. 3C). These patterns are consistent with horizontal gene transfer from non-viral hosts. In contrast, other taxonomically distributed protein clusters such as cluster 56 (encoding parvovirus Rep proteins with homologs in some human herpesviruses) and cluster 735 (encoding a hemagglutinin lineage present in baculoviruses and some orthomyxoviruses) display a monophyletic taxonomic distribution consistent with horizontal gene transfer between viruses (Supplementary Fig. 3D, E). These data suggest that many protein clusters that contain proteins from viruses of different genome types arise from horizontal gene transfer, both from viral and non-viral sources.

### Structural alignments identify shared functional domains

We constructed protein clusters with a strict 70% coverage requirement, leaving open the possibility that individual domains can be identified through structure comparison^3^. We reasoned that protein domains present within multiple protein clusters may have particular biological importance. We used DALI to conduct all-by-all alignments of the representatives of all protein clusters having more than one member. This revealed substantial protein similarity with many alignments having Z scores greater than 8, indicating high confidence of structural homology^25^ (Supplementary Fig. 4A). Protein clusters ultimately fall into a network of shared domains (Supplementary Fig. 4B). Here, distinct domains are often shared across protein clusters in context with various combinations of other domains, which can be seen with domains involved in interaction with the cytoskeleton (Supplementary Fig. 4C) and in metabolism (Supplementary Fig. 4D,E) in eukaryotic viruses and phage.

### Structural homology reveals phosphodiesterases that degrade 2’3’ cGAMP

Many aspects of eukaryotic and prokaryotic immunity have a shared origin^41^. One set of related pathways are the mammalian cGAS-STING and OAS pathways and prokaryotic Cyclic-oligonucleotide-based anti-phage signaling systems (CBASS). In both cases, a protein sensor detects a viral cue and generates a nucleotide second messenger, which activates a downstream antiviral effector (Fig. 4A). In the case of the cGAS (cyclic GMP-AMP synthase) pathway, cGAS recognizes cytoplasmic double-stranded DNA and generates 2’3’ cyclic GMP-AMP (2’3’ cGAMP). Many cGAS/DncV-like nucleotidyltransferases (CD-NTases) in prokaryotic CBASS’ make a similar second messenger, 3’3’ cGAMP, in response to viral cues^42^. In contrast, OAS (oligoadenylate synthase) recognizes double-stranded RNA and generates linear 2’5’ oligoadenylates (2’5’ OA)^43,44^. In prokaryotes, phage T4 encodes the ligT-like PDE anti-CBASS protein 1 (Acb1), which degrades 3’3’ cGAMP and a variety of other cyclic nucleotide substrates including 2’3’ cGAMP^45^.

**Fig. 4:**
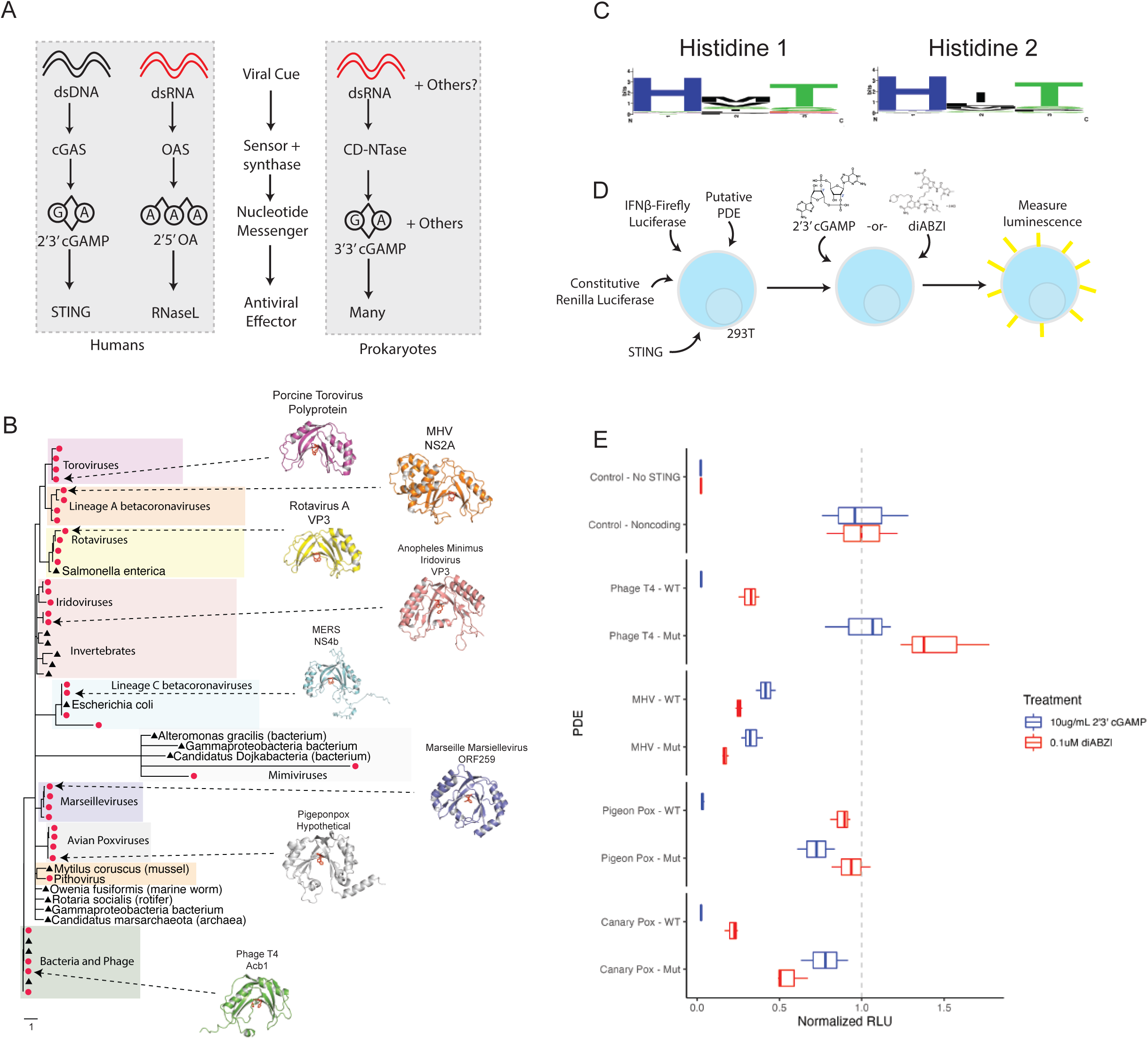
LigT-like phophodiesterases are frequently used to subvert host immunity. (**A**) Some innate immune pathways in eukaryotes and prokaryotes rely on a viral synthase sensor that detects virus-associated molecular patterns such as dsDNA or dsRNA and generates a nucleotide second messenger that stimulates an antiviral effector. This includes the human cGAS and OAS pathways, as well as prokaryotic CBASS pathways. (**B**) A phylogenetic tree showing the polyphyletic lineages of ligT-like PDEs. A structural search was used to identify viral proteins with a conserved ligT-like fold, and close viral and non-viral members of each were used to construct a phylogenetic tree. Shaded boxes indicate a lineage, with red nodes indicating viral proteins and black nodes indicating non-viral proteins. The red residues in each protein structure are the conserved catalytic histidines. (**C**) A sequence logo showing conservation of the catalytic histidines in viral ligT-like PDEs despite exceptionally low sequence identity overall. (**D**) Outline of the experimental system to assess degradation of 2’3’ cGAMP by viral ligT-like PDEs. HEK 293T cells were transfected with constructs encoding STING, firefly luciferase driven by an IFN-beta promoter, a constitutive renilla luciferase, and a transgene. After 5 hours, cells were treated with 10ug/mL 2’3’ cGAMP or 0.1uM diABZI (a non-cyclic dinucleotide STING agonist). Around 24 hours after the first transfection, luminescence of the firefly and renilla luciferases were measured. (**E**) LigT-like PDEs from avian poxviruses are active against 2’3’ cGAMP. X axis indicates the relative RLU, with all diABZI-treated conditions being normalized to the diABZI RLU of cells transfected with a noncoding transgene and all 2’3’ cGAMP-treated conditions being normalized to the 2’3’ cGAMP RLU of cells transfected with a noncoding transgene. RLUs were initially normalized as (firefly RLU)/(renilla RLU). The Y axis indicates the transgene, with the color of each box indicating the treatment (cGAMP or diABZI). “Mut” indicates the transgene contains H→A mutations of both catalytic histidines.

In eukaryotes, several RNA viruses encode PDEs that degrade 2’5’ OA^46^. Interestingly, we find that these PDEs have a ligT-like fold similar to Acb1. Given the conserved use of ligT-like PDEs in viral anti-immunity, we investigated their distribution and phylogeny. Structural searches revealed many different branches of ligT-like PDEs are present in eukaryotic viruses (Fig. 4B). Notably, there are multiple independent branches of ligT-like PDEs in RNA viruses. Linage A betacoronaviruses and Toroviruses share a clade of PDEs that is similar to the PDEs present in Rotaviruses. Surprisingly, lineage C betacoronaviruses contain a distinct branch of PDEs (Fig. 4A)^47^. This suggests that there were two independent PDE acquisition events within the betacoronavirus genus, showing the strong selective pressure for betacoronaviruses to evade the OAS pathway. We find that some large DNA viruses also contain ligT-like PDEs. Despite the extreme amino acid variability across the ligT-like PDE tree there is near-universal conservation of the two catalytic histidines (Fig. 4C), with the exception of the Mimivirus ligT-like branch.

The presence of ligT-like PDEs in large DNA viruses raises the question of whether they have an anti-immune function. While the RNA-sensing OAS pathway is commonly targeted by ligT-like PDEs of RNA viruses, there is likely less pressure for large DNA viruses to target OAS. Thus, we tested whether ligT-like PDEs encoded by large DNA viruses have activity against 2’3’ cGAMP. To address this question, we generated a synthetic STING circuit in 293T cells^48,49^ (Fig. 4D). Here, STING can be activated by treatment with 2’3’ cGAMP or the non-nucleotide STING agonist diABZI^50^, which will lead to expression of firefly luciferase in a STING-dependent manner. We expect that a viral PDE that targets 2’3’ cGAMP should be able to inhibit 2’3’ cGAMP- but not diABZI-mediated STING activity. Testing representative PDEs from each branch revealed that while PDEs encoded by RNA viruses and other large DNA viruses have only limited activity against 2’3’ cGAMP, PDEs encoded by avian poxviruses have very potent activity against 2’3’ cGAMP (Supplementary Fig. 5A). We found that the ligT-like PDEs encoded by Pigeonpox and Canarypox very potently restrict STING signaling stimulated by 2’3’ cGAMP but have limited activity against diABZI-mediated STING signaling (Fig. 4E). Furthermore, mutation of the catalytic histidines substantially reduces activity (Fig. 4E). Avian poxvirues are notable for their lack of Poxin^51^, the other 2’3’ cGAMP phosphodiesterase encoded by poxviruses, showing the strong selective pressure for poxviruses to evade cGAS-STING immunity. Altogether, we have leveraged structure homology to discover a novel mechanism of 2’3’ cGAMP degradation by eukaryotic viruses and find that cGAMP targeting by ligT-like PDEs is a pan-viral mechanism of anti-immunity.

## Discussion

Viruses have yielded fundamental insights into basic molecular biology. Here, we cluster viral proteins and use structural alignments to gain functional insights missing with prior approaches. We have raised testable hypotheses about the function of proteins present in human pathogens, and provided a resource for studying viral protein structures at scale. Expanding databases of structures and predicted structures will continue to enable functional inference. This is important not only from a fundamental biology perspective, but also in light of the continual emergence of novel viruses with pandemic potential. Structural homology both with other viral proteins and with host proteins can offer functional insights and provide insight into the origin and evolution of viral proteins.

A caveat of our study is the use of a stringent 70% coverage threshold. This means that some proteins with similar function but differences in domain configuration will be split into separate protein clusters, underestimating their taxonomic diversity. However, we find that structure alignment-based domain identification can identify shared structural repeats and enable comparison across protein structures. Both structure prediction and alignment have fundamental limitations. These structures are predictions, whose quality can vary and can be influenced by the depth of the multisequence alignment used for prediction. Structural alignments in turn can be affected by the arbitrary positioning of mobile domains.

Protein structure is especially informative in cases of evolutionary distance. One impactful area is the concept of conserved viral anti-immunity. Emerging evidence on the common origin of some bacterial and eukaryotic immune systems raises the potential for conserved anti-immunity systems in eukaryotic viruses and phage. This is illustrated by ligT-like PDEs, which have been adapted multiple times by phage and eukaryotic viruses to evade innate immunity. This also illustrates the flexibility of core protein folds, as this conserved fold can be adapted to cleave distinct immune second messengers depending on the pathway being targeted. Altogether, our study lays the foundation for characterization of viral protein evolution and function across the virome.

## Methods

### Preparation of protein sequences

Protein sequences for eukaryotic viruses present in RefSeq^52^ were collected through the NCBI Viruses portal (https://www.ncbi.nlm.nih.gov/labs/virus) in July 2022. GenPept files were downloaded for viruses that were annotated by NCBI to have a eukaryotic host. Because not all viruses have a host labeled by NCBI, GenPept files of human-infecting viruses annotated by ViralZone (https://viralzone.expasy.org/678) were also downloaded. Finally, proteins from all coronaviruses present in RefSeq, regardless of NCBI-labeled host, were downloaded.

Each GenPept file was processed such that polyproteins with defined “mature peptide” fields produced separate protein sequences for each mature peptide. GenPept files without a mature peptide field were output as full amino acid sequences. These processing steps are present in the vpSAT github directory (https://github.com/jnoms/vpSAT) in the process_gbks.py file. Proteins larger than 1500 residues, or in some cases 1000 residues, were excluded. This amounted to only 1,706 excluded proteins.

### Structure prediction

Multisequence alignments (MSA) were generated with MMseqs2 release version b0b8e85f3b8437c10a666e3ea35c78c0ad0d7ec2. To increase MSA generation speed, the RefSeq virus protein database (downloaded on June 6, 2022) was used as the target database for MSA generation. Structures were predicted with Colabfold^20^, downloaded June 22, 2022. The majority of samples used three recycles, three models, stop_at_score=70, and stop_at_score_below=40. MMseqs2 and Colabfold_batch were run with a Nextflow^53^ pipeline, and all parameters used can be found here: https://github.com/jnoms/vpSAT. Information on all viruses and structures included in this manuscript is present in **Supplementary Table 1**.

### Protein cluster generation

All proteins were initially clustered with MMseqs2, with a requirement of at least 20% sequence identity and 70% query and target coverage. MMseqs2 cluster mode 0 was used, meaning that many but not all pairs of aligned proteins are placed into the same sequence cluster. Predicted structures for each sequence cluster representative were subjected to an all-by-all alignment using Foldseek^22^, requiring the alignment to consist of at least 70% query and target coverage and an alignment *E*-value less than 0.001. The resultant structural alignment file was then filtered using SAT aln_filter to keep alignments with a TMscore of at least 0.4. Clusters were generated from this alignment file using SAT aln_cluster in a similar manner as Foldseek cluster mode 1, wherein all query-target pairs are assigned to the same cluster. Cluster information from sequence and structure clustering were merged using SAT aln_expand_clusters. Taxonomic counts information was generated using SAT aln_taxa_counts, producing a “tidy” table for each cluster_ID with the number of members of each taxon at multiple taxonomy levels. Taxonomy information was also added directly to the merged cluster file using SAT aln_add_taxonomy.

### Cluster purity analysis

To determine the structural consistency of the clusters, all clusters with at least 100 members were selected for analysis. DALI was used to align the cluster representative with each cluster member. Clusters whose members were on average smaller than 150 residues were excluded. This led to the analysis of 49 clusters. Cluster members that failed to align to their representative were assigned a Z value of 0. For each cluster, the average Z score between the representative and each member was determined and plotted. All scripts used to run DALI can be found in vpSAT’s dali_format_inputs.sh and dali.sh files. Dalilite version 5 was used. DALI output files were parsed into a tabular format using SAT’s aln_parse_dali.

### Phylogenetics

To generate phylogenetic trees, one or more sequence cluster representatives (or, in the case of phosphodiesterases, one representative from each clade) were searched against the NCBI non-redundant database using web BLASTP. The top three viral and top three non-viral hits were collected. Multiple sequence alignments were generated using MAFFT^54^ FFT-NS-i x1000, gap open penalty of 1.53, offset value 0.123, implemented in Geneious. Phylogenetic trees were generated using the Geneious Tree Builder with a Jukes-Cantor genetic distance model.

### Structural alignments against the Alphafold databases

In Fig. 1I, Foldseek was used to align a protein representative from every viral protein cluster against 2.3 million protein cluster representatives from the Alphafold Database^3^. For Fig. 3, all 67,715 viral protein structures were searched against the pre-made Foldseek databases of the original release of the Alphafold database, consisting of proteins from 48 organisms and including members of the bacterial, eukaryote, and archaeal superkingdoms. For this search, the full Alphafold database of over 200M structures was not used because it contains many viral proteins misannotated as non-viral proteins (these misannotations reflect errors in Uniprot metadata). Alignments were filtered to keep only those with a minimum TMscore of 0.4 and an *E*-value of less than 0.001.

**DALI alignments of specific non-viral proteins against the viral protein database** Following Foldseek alignments against the Alphafold database, specific hits of interest (e.g. ENT4) were selected. These structures were downloaded and imported to the DALI database format using vpSAT’s dali_format_inputs.sh. They were then aligned against the full viral protein structure database using vpSAT’s dali.sh, which lists all parameters. Dalilite version 5 was used. DALI output files were parsed into a tabular format using SAT’s aln_parse_dali.

### Identification of annotated protein sequence clusters

Each protein in the database was searched against the Pfam^29^, CDD^30^, and TIGRFAM^31^ databases using InterProScan^28^. A sequence cluster was considered “annotated” if greater than 25% of members had any InterProScan alignment, and was considered “unannotated” if otherwise. RMSDs present in Fig. 2 were calculated using DALI.

### DALI alignments to identify shared domains

This analysis used the structure representatives from structures with at least 2 members, resulting in 5.7K cluster representatives. Structures from these representatives were imported to the DALI database format using vpSAT’s dali_format_inputs.sh. To compare eukaryotic virus protein cluster representatives, an all-by-all alignment was conducted using vpSAT’s dali.sh, which lists all parameters.

Dalilite version 5 was used. DALI output files were parsed into a tabular format using SAT’s aln_parse_dali. All DALI alignments were filtered for an alignment length of at least 120, and for a Z score greater than or equal to (alignment length/10) - 4.

### Data analysis and plotting

All analysis and plotting used R version 4.0.3. The genome type and average genome size were determined from information downloaded from NCBI’s Virus portal (https://www.ncbi.nlm.nih.gov/labs/virus/vssi/#/).

### Phosphodiesterase Activity Assay

293T cells were seeded into 96-well plates at 20,000 cells per well. The day after plating, cells was transfected with 15ng STING (pMSCV-hygro-STING R232, Addgene 102608), 20ng encoding firefly luciferase driven by an IFN-beta promoter (IFN-Beta_pGL3, Addgene 102597), 5ng of renilla luciferase (pRL-TK - Promega E2241), and 20ng of each putative PDE using the Mirus TransITX2 transfection reagent. After at least 4 hours, cells were treated with 0.1uM diABZI (Invivogen) or transfected with 10ug/mL 2’3’ cGAMP (Invivogen) using TransITX2. The next day, firefly and renilla luciferase were measured using the Promega Dual-Glo luciferase assay system. Three wells were transfected per condition, and experiments are representative of at least 2 independent experiments.

### Data and Code Availability

Code used for upstream processing is present in the vpSAT Github repository (https://github.com/jnoms/vpSAT/tree/main). This includes scripts required for most computational steps. A workflow is available that shows all main processing steps: https://github.com/jnoms/vpSAT/blob/main/manuscript_code/2024-01-04/analysis_workflow.ipynb.

The SAT python package can be downloaded as instructed on the SAT Github repository (https://github.com/jnoms/SAT/tree/main).

All plotting and analysis scripts are available as Quarto documents: https://github.com/jnoms/vpSAT/blob/main/manuscript_code/2024-01-04/.

The vpSAT and SAT github repositories are private during review, but all code can be accessed in the Zenodo repository listed below, along with all intermediate data necessary to reproduce the figures: https://doi.org/10.5281/zenodo.10460192

Finally, all structures are available on Zenodo: https://doi.org/10.5281/zenodo.10291581

## Acknowledgements

J.A.D. is an Investigator of the Howard Hughes Medical Institute. Portions of this research were conducted on the Wynton Cluster at UCSF, supported by UCSF IT. We would like to thank members of the Doudna lab for their support and helpful feedback.

## Author Contributions

Conceptualization - J.N

Methodology - J.N., J.A.D.

Software - J.N.

Validation - J.N.

Formal Analysis - J.N.

Investigation - J.N., N.P.

Resources - J.N., J.A.D.

Data Curation - J.N.

Writing, Original Draft - J.N.

Writing, Review and Editing - J.N., N.P., J.A.D.

Visualization - J.N.

Supervision - J.A.D.

Project Administration - J.N., J.A.D.

Funding Acquisition - J.A.D.

---

**Supplementary Fig. 1:**
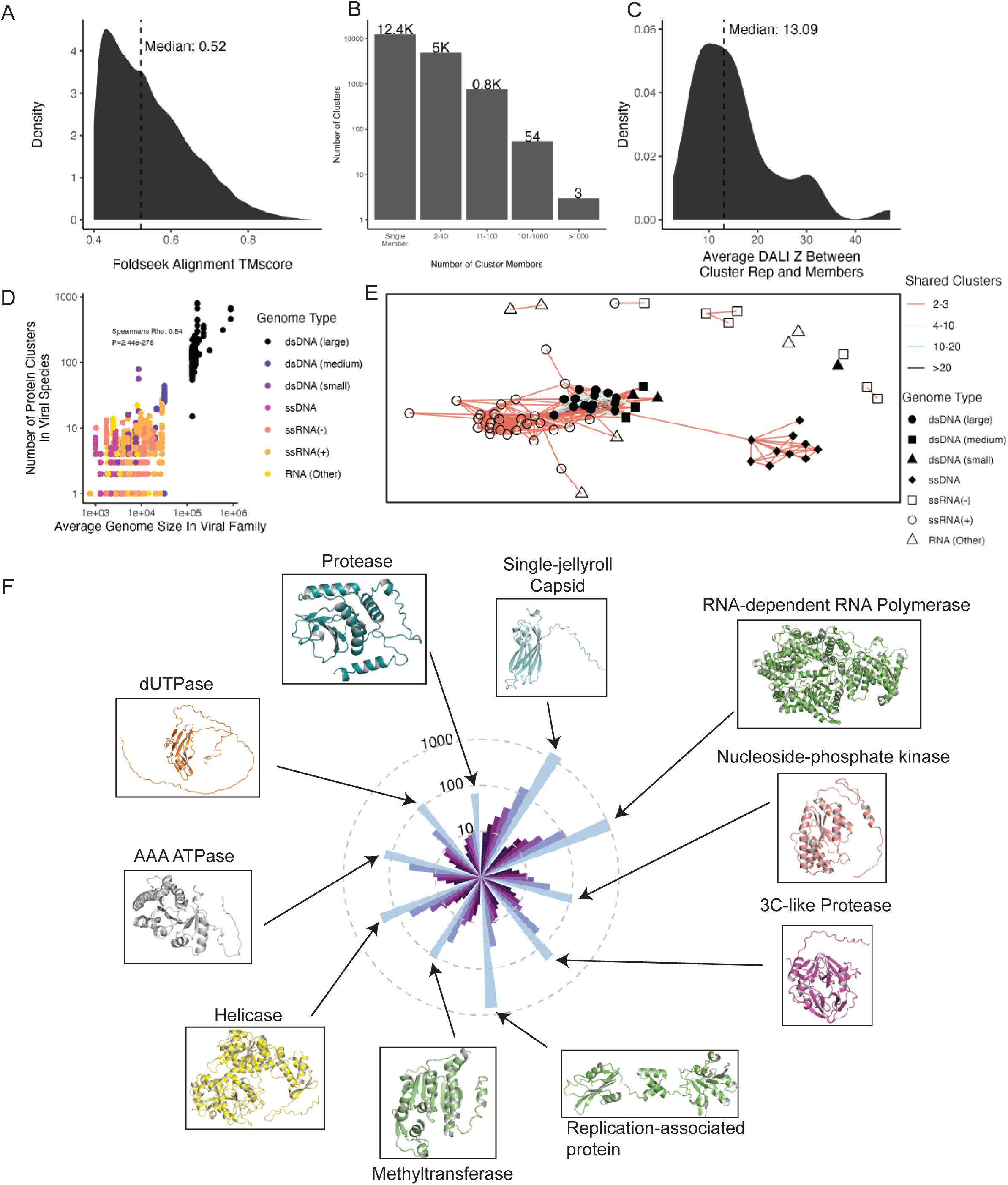
Distribution of protein clusters across viral families. (**A**) Foldseek was used to align all virus sequence cluster representatives against one another, and alignments with a TMscore below 0.4 were removed. This plot shows the distribution of alignment TMscores, with the X axis indicating the TMscore and the Y axis indicating the density (or “proportion”) or alignments with each TMscore. (**B**) The distribution of proteins amongst sequence clusters. The X axis indicates the size of each cluster, while the Y axis indicates the number of clusters of that size. (**C**) For each protein cluster with at least 100 members, the cluster representative was aligned with DALI against all cluster members. Clusters that contained members with an average length of 150 residues or less were excluded, and members that did not align to the representative were assigned a Z score of 0. The distribution of average Z scores for each cluster is plotted, with the median cluster-averaged indicated. X axis indicates the DALI Z score for each cluster, while the Y axis indicates the density (or proportion) of clusters with each average DALI Z score. (**D**) Relationship between the number of protein clusters encoded by a viral species (Y axis) and the average genome size of its family (X axis). Each dot is a viral species, and colors indicate the genome type. The spearman’s Rho is 0.54, with a P value of 2.44e-278, indicating a strong correlation. (**E**) Each node represents a single viral family, with the shape and color indicating the genome type of that family. The color of edges between the nodes indicates the number of shared protein clusters between each pair of families. Only those family-family pairs with at least 2 shared protein clusters are plotted. (**F**) Protein clusters were ordered by their phylogenetic diversity of their members (e.g. # phyla > # classes > # orders >… # species) and the top 10 clusters were plotted. Bars are colored based and ordered on decreasing taxonomic level, with phyla as dark blue on the far left and species as bright blue on the far right of each stack.

**Supplementary Fig. 2:**
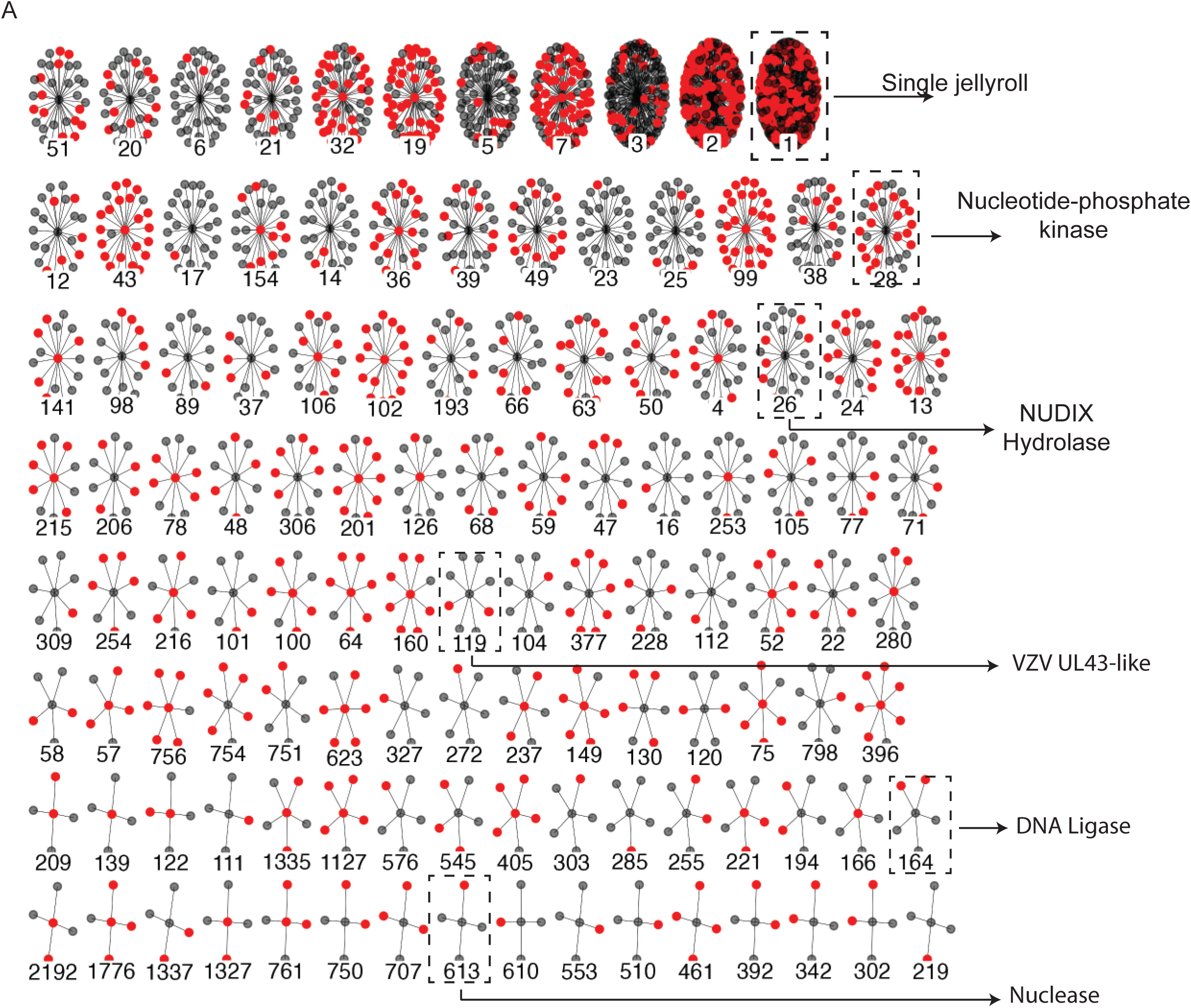
Many unannotated proteins have structural homlogy to annotated protein clusters. (**A**) Many protein clusters contain a mix of annotated and unannotated sequence clusters. Each “wheel” of nodes indicates a protein cluster, with individual nodes representing individual sequence clusters. Each sequence cluster node is colored based on if it is annotated (gray) or unannotated (red). All protein clusters with at least one annotated and one unannotated protein cluster are shown.

**Supplementary Fig. 3:**
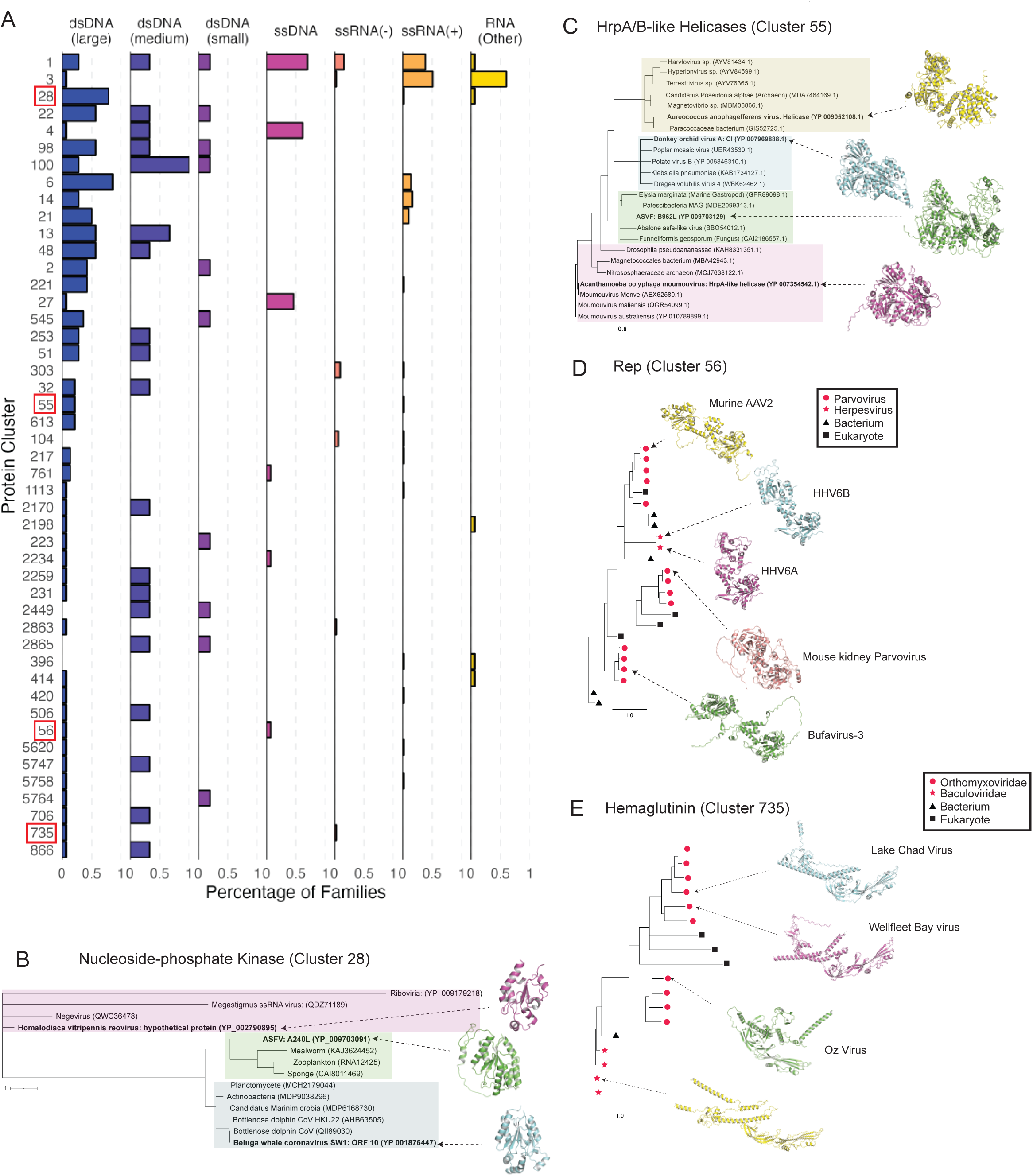
Horizontal gene transfer drives the emergence of taxonomically-diverse protein clusters. (**A**) Protein clusters were ranked as follows: 1) by the number of genome types of viral species that encode cluster members, followed by 2) the number of viral families that encode cluster members. The top 50 protein clusters by this metric were included in the plot. Each row is a protein cluster (with the number indicating the protein cluster ID). The X axis indicates the percentage of viral families of each genome type that contain a viral species that encodes a member of the protein cluster. (**B**) A polyphyletic protein cluster of a nucleotide-phosphate kinase fold. The closest viral and non-viral sequence homologues of three viral sequence representatives were used to generate a phylogenetic tree of protein cluster 28. Shaded boxes indicate distinct phylogenetic branches. The scale bar indicates substitutions per site. (**C**) A polyphyletic protein cluster of HrpA/B-like helicases. The closest viral and non-viral sequence homologues of four viral sequence representatives were used to generate a phylogenetic tree of protein cluster 55. Shaded boxes indicate distinct phylogenetic branches. The scale bar indicates substitutions per site. (**D**) A monophyletic protein cluster of Rep-like proteins shows sequence similarity between Parvovirus Rep proteins and a Rep-like protein in HHV6A and HHV6B. The closest viral and non-viral sequence homologues of five members of protein cluster 56 were used to generate a phylogenetic tree. Red nodes indicate viral sequences, with circles corresponding to parvoviruses and stars indicating herpesviruses. Black nodes indicate non-viral sequences, with triangles indicating bacteria and squares indicating eukaryotes. The scale bar indicates substitutions per site. (**E**) A monophyletic protein cluster of Hemaglutinin-like proteins shows sequence similarity between a clade of orthomyxovirus and baculovirus hemaglutinins. Red nodes indicate viral sequences, with circles corresponding to orthomyxoviruses and stars indicating baculoviruses. Black nodes indicate non-viral sequences, with triangles indicating bacteria and squares indicating eukaryotes. The scale bar indicates substitutions per site.

**Supplementary Fig. 4:**
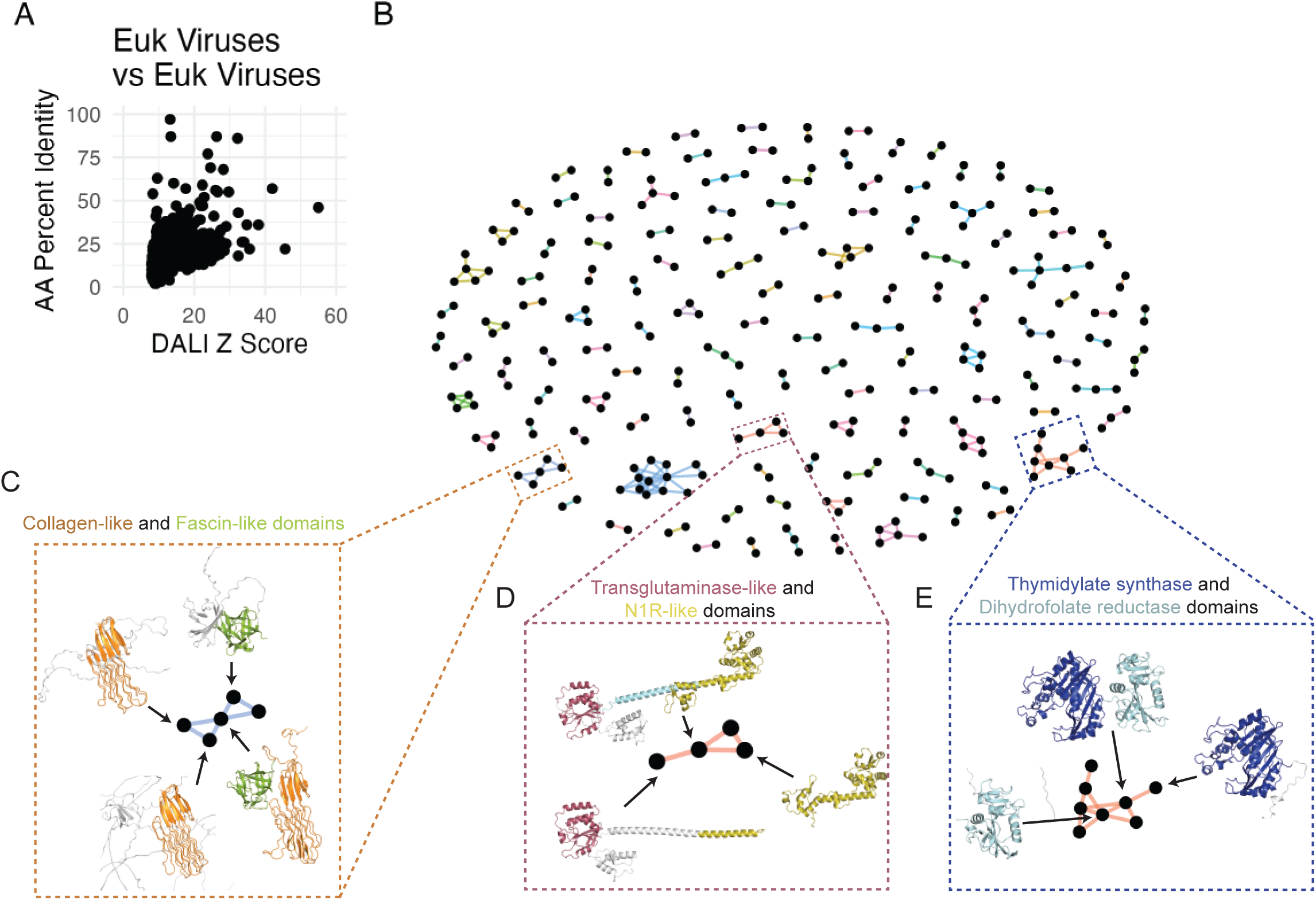
Shared domains across eukaryotic virus protein clusters. (**A**) All-by-all structural alignments of representative structures from the 5,770 protein clusters with more than one member. Each dot indicates a single alignment, with the Y axis indicating the fraction of amino acid identity and the X axis indicating DALI Z-score. The color of each dot indicates the alignment coverage (between 0 and 1), where coverage is (length of alignment)/(length of query or target, whichever is longest). (**B**) Protein clusters tend to share protein domains. Each node indicates a protein cluster, and edges between protein clusters indicates there is a DALI alignment between them. Only alignments with a Z score of at least 15 are plotted. The boxes indicate cluster representatives highlighted in subsequent panels. (**C**) Frequent reuse of structural/cytoskeleton-related domains. Protein clusters with collagen-like domains (orange) and fascin-like domains (green) are highlighted. (**D**) Multiple combinations of domains with the same viral genus. Diverse combinations of transglutaminase-like domains (purple) and N1R-like domains (green) from entomopoxvirus proteins are highlighted. (**E**) Frequent reuse of protein domains involved in metabolism. Various combinations of thymidylate synthase (dark blue) and dihydrofolate reductase (light blue) domains in eukaryotic protein clusters are highlighted.

**Supplementary Fig. 5:**
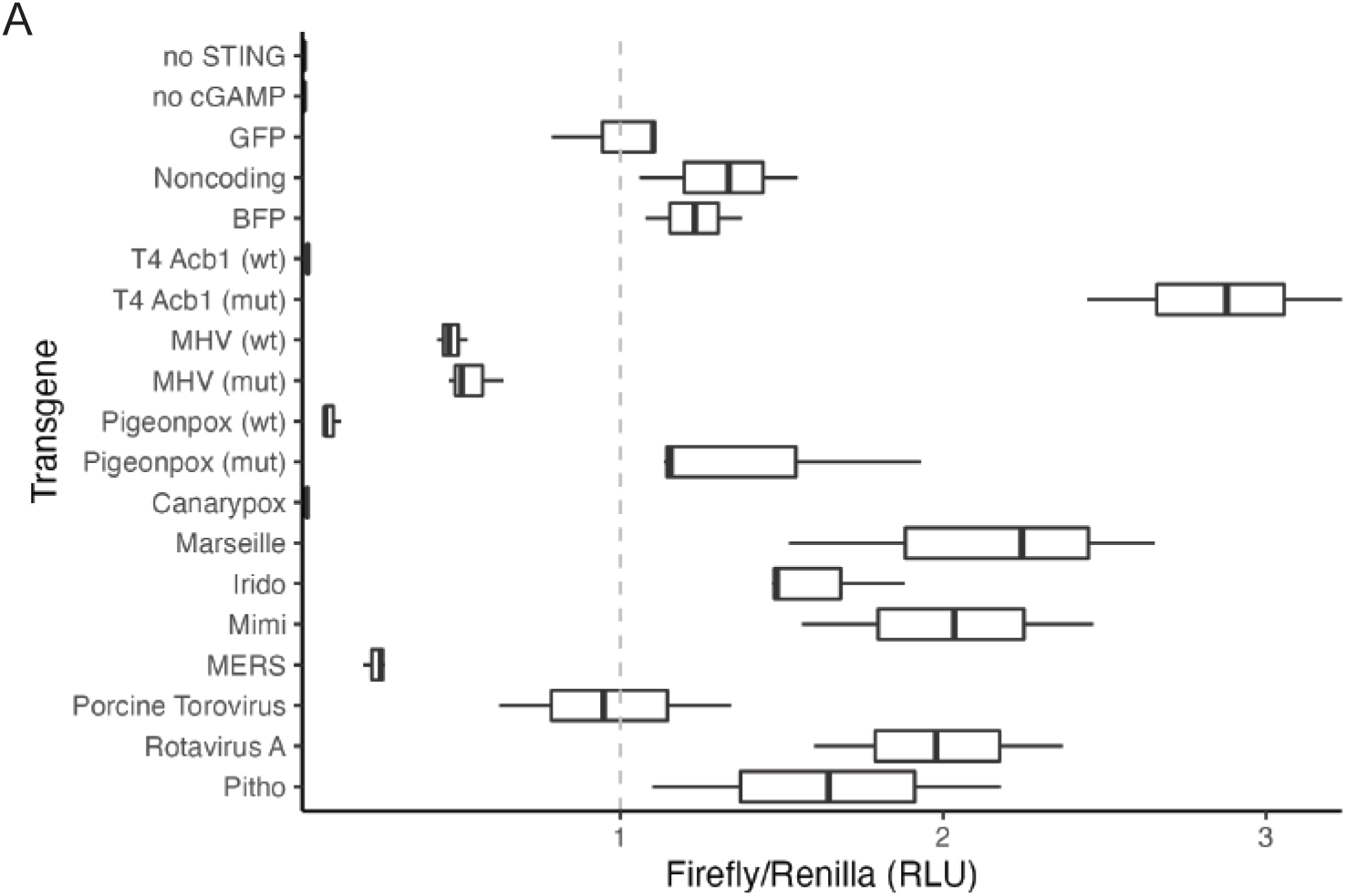
Activity of viral PDEs against 2’3’ cGAMP. (**A**) X axis indicates the relative RLU normalized to the GFP transgene condition. ”Mut” indicates the transgene contains H→A mutations of both catalytic histidines.

